# Improving Biological Sequence Prediction with AlphaFold2 Representation

**DOI:** 10.64898/2026.04.26.720550

**Authors:** Zhiqian Jiang, Canh Hao Nguyen, Hiroshi Mamitsuka

## Abstract

**Motivation:** Accurate prediction of functional sites from primary sequences is essential for elucidating biological mechanisms and advancing rational drug design. However, traditional sequence-based features are inherently unable to capture complex structural protein contexts. Recently, AlphaFold2 (AF2) revolutionized protein structure prediction, raising expectations of AF2 to serve as a feature extractor providing structure-rich representation, which can be useful for sequence-based prediction, particularly for unknown sequences.

**Results:** We present a novel feature-engineering paradigm that leverages a high-dimensional latent representation matrix (of *L × D*, where *L* is the sequence length and *D* is the feature dimension size) extracted directly from the AF2 Evoformer module. We systematically evaluated the AF2 representation, comparing with conventional sequence-based features, such as hidden Markov model profiles, using a variety of machine learning models, on two structurally contrasting tasks, calpain cleavage site and nucleic-acid-binding site prediction. The AF2 representation outperformed conventional sequence-based features clearly and entirely, particularly for targets with low sequence homology to training data. Furthermore, interpretability analyses, using SHapley Additive exPlanations (SHAP) and Uniform MAnifoldapproximation and Projection (UMAP), showed more details behind the performance advantage of AF2 representation through feature importance ranking and visualization. Overall, these empirical results confirmed that AF2 representation could effectively bridge the sequence-to-structure gap as a feature input for sequence prediction, without increasing heavy computational burden.

**Availability and implementation:** Source code, pre-trained models, and datasets are freely available to non-commercial users at https://github.com/Lili-irtyd/Improve-biological-sequences-prediction-by-AlphaFold2.

## Introduction

Identifying functional sites, such as enzymatic cleavage regions, ligand-binding pockets, and interaction interfaces, from primary amino acid sequences is a foundational endeavor in structural bioinformatics. Accurate finding of these sites in sequences is pivotal for elucidating protein functions, understanding cellular mechanisms, and accelerating rational drug design and protein engineering. Traditionally, functional site prediction has relied on accurate, experimental structural biology, which is, however, resource-intensive and unable to keep pace with the exponential growth of sequencing data. Consequently, computational approaches have become indispensable for bridging the sequence-to-function gap.

Computational methods for protein sequence analysis have undergone a significant evolution. Early approaches heavily depended on statistical models and traditional machine learning (ML) algorithms, such as Position-Specific Scoring Matrices (PSSMs) and Hidden Markov Models (HMMs), to capture evolutionary conservation. Recently, Protein Language Models (PLMs, e.g., ESM-2) have revolutionized the field by deeply learning sequence syntax and co-evolutionary patterns from massive sequence databases [1]. Concurrently, breakthrough architectural innovations like AlphaFold2 (AF2) [2] have largely solved the protein folding problem. By leveraging the explicit integration of pairwise physical constraints and spatial attention in the Evoformer module, AF2 achieves atom-level accuracy in protein three-dimensional (3D) structure prediction.

Despite these remarkable advancements, efficiently extracting structurally aware features for input sequences to improve predictions for downstream tasks remains unresolved. While PLMs excel at sequence-level modeling, they inherently lack explicit spatial geometry and often struggle to capture long-range 3D dependencies. On the other hand, generating the final 3D coordinates by AF2 requires a massive computational burden, particularly for flexible or intrinsically disordered regions. In this context, an intermediate outcome in AF2, i.e. the latent vector (or matrix) representation, which hereafter we call **single representation**, of the Evoformer remains underexplored. More practically, in this study, this representation is a continuous residue-level feature matrix of *L × D*, where *L* denotes the protein sequence length and *D* corresponds to the size of the latent channel dimension (384 in our experiments). Generated prior to the 3D coordinate generation module, this dense feature matrix must contain information on both multiple sequence alignment (MSA) and implicit 3D spatial semantics. We hypothesize that this matrix serves as a compact, structurally-aware information that provides a significantly richer context than simpler sequence-based features, allowing to bypass the need for explicit 3D coordinate generation.

It is essential to rigorously validate the universality and robust generalizability of this latent representation, across tasks with fundamentally different structural and biological dependencies. To this end, as validation tasks for single representation from AF2, we deliberately selected two predictive downstream tasks situated at opposite ends of the structural dependency spectrum:

- **Calpain cleavage site prediction:** This task of identifying calpain cleavage sites in a given input sequence is considered to be a process governed by local and flexibly accessible motifs [3, 4]. Calpains recognize substrate sequences that are often located in intrinsically disordered regions [5, 6]. This task examines the capability of the representation, regarding capturing subtle, short-range sequence determinants and local conformational flexibility, excluding the rigid secondary structures.
- **Nucleic-acid-binding site prediction:** In contrast, recognizing DNA (or RNA) binding sites in a given sequence demands the perception of rigid, global spatial clustering and long-range electrostatic surface properties [7, 8]. Binding residues are often far apart in the primary sequence but near together in the 3D space. This task examines the capacity of the representation regarding the inherent encoding of long-range, complex steric information.

By evaluating the AF2 representation across these two contrasting scenarios, i.e. one driven by local sequence flexibility and the other by global structural rigidity, we aim to provide a comprehensive assessment of the broad versatility as a feature extractor. In this paper, we systematically extract and integrate the single representation of the AF2 Evoformer into a diverse array of predictive ML architectures, ranging from Support Vector Machines (SVMs) to Graph Neural Networks (GNNs), to evaluate the efficacy as a standalone feature for functional site prediction.

The main contributions of this work are as follows:

- We present a novel feature-engineering paradigm that utilizes the latent single representation from AlphaFold2, effectively incorporating implicit 3D spatial semantics without the computational burden of full 3D coordinate generation.
- We demonstrate the broad universality of this representation by successfully applying to two structurally contrasting tasks: local, flexible calpain cleavage site prediction and global, rigid nucleic-acid-binding site prediction.
- We systematically validate that implicit, structural representation by AF2 keeps high generalizability across various ML models, e.g. SVMs and GNNs.
- Empirical validation results reveal that single representation by AF2 significantly outperformed conventional sequence-based baselines in a variety of settings of the two downstream tasks, offering a robust alternative when experimental or highly accurate 3D structures are unavailable.

### Methodology: Generating “AF2 Feature” by Repurposing AlphaFold2 for Feature Extraction

AlphaFold2 (AF2) is designed as an end-to-end three-dimensional (3D) structure predictor, to output the final 3D atomic coordinates. We retrieved features for sequences from AF2 by repurposing AF2 into a high-throughput feature extractor, focusing specifically on the latent vector (or matrix) representation (which is hereafter called **single representation**), implicitly generated by the Evoformer module of AF2. This high-dimensional vector must contain the information on both multiple sequence alignment (MSA) and implicit 3D spatial semantics prior to 3D coordinate generation.

To efficiently extract this dense, residue-level embedding matrix as a standalone feature for downstream analysis, we systematically modified the AF2 inference pipeline. These modifications isolate the core representation, prevent structural bias, and significantly reduce computational load.

#### Isolating the Core Representation

To extract the intermediate single representation, we altered the core model configuration by setting `return_representations=True` within the `AlphaFold (hk.Module)` class. This forces the model to bypass the standard coordinate output and instead return a dictionary (`result_model_*.pkl`) containing intermediate representation tensors.

We specifically isolated the single representation by accessing the `[‘representations’][‘single’]` key path. Furthermore, we modified the return values across the pipeline to ensure that only this specific matrix (single representation) is returned, discarding the rest of the dictionary to minimize memory requirements.

#### Disabling Iterative Recycling (num_recycle=0)

AF2 natively employs a recycling mechanism, passing outputs back through the network multiple times to refine structural predictions. We disabled this iteration by setting `num_recycle = 0` to minimize the number of ensemble models, considering the balance between feature refinement and computational cost. That is, this iterative process is an optimization, in terms of structural feature refinement, of the output, which might already have sufficient MSA and structure information, even by the result of a single run. Thus, this iteration might not improve the single representation much, for the increased computational cost by multiple runs. In fact, bypassing this iteration drastically reduces the computational load and accelerates inference.

#### Bypassing Auxiliary and Structural Modules

Our target is solely the output of the Evoformer, and downstream modules in AF2 are irrelevant and thus can be functionally disabled or skipped in `modules.py` and `model.py`, particularly for computational efficiency. Overall, our strategy of “keep and discard” was as follows:

- **Components retained:** We preserved the core `Embeddings_AndEvoformer` module (responsible for generating the initial sequence embeddings and evolutionary information) and `ensemble_representations` (to combine paths and ensure stability).
- **Auxiliary heads removed:** We bypassed computationally intensive auxiliary heads, including `MaskedMsaHead` and `DistogramHead`, since we do not need to predict MSA masks or distance distributions. We also removed the `compute_loss` module, which is unnecessary during inference.
- **Confidence scoring skipped:** Computations for the predicted local distance difference test (pLDDT) and predicted aligned error (PAE) scores, which are both designed to evaluate the confidence of 3D atomic predictions, were removed from `run_alphafold.py` to save resources.

#### Pipeline Optimization for High-Throughput Inference

To transform the modified AF2 script into a high-throughput feature extraction tool, we cleaned the execution environment by removing the benchmark-related code and the verbose log output. This minimizes I/O bottlenecks, resulting in a cleaner execution stream. Finally, to handle large-scale datasets, we wrapped the modified pipeline in a custom batch-processing script (`process_fasta_files.py`). This script automatically iterates over directories of FASTA files, feeding them sequentially into the streamlined AF2 pipeline to generate and save the intermediate single representation matrices in a continuous, automated loop.

#### Sequence Length Limitation by Memory Consumption of AF2

A known computational constraint of the AF2 architecture is, because of the high GPU memory consumption, the inherent limitation on long sequence processing. Due to this hardware limitation, sequences must be restricted by a specific cut-off length to fit within GPU VRAM limitations. Consequently, all proteins used in two downstream tasks in this work were strictly selected or truncated to fall below a cut-off of 1,000 residues to ensure successful feature extraction.

### Downstream Tasks

#### 1. Calpain Cleavage Site Prediction

##### Benchmark Datasets

We used proteins of the dataset in [6] under the condition that the length of a protein sequence is less than 1,000 residues, resulting in 48 proteins. For feature extraction, we used a sliding window strategy, defining each cleavage site as the central residue with 15 flanking residues on each side, forming a 31 amino acid window (*L* = 31). For generating AF2 single representation, full-length sequences were used to preserve their global structures; the 31-residue windows were sliced from the resulting feature matrices of the entire sequences. To strictly prevent data leakage and ensure a rigorous evaluation of generalization, the 48 proteins were partitioned into a training set (80%) and an independent test set (20%) such that no highly homologous sequences (e.g., *>* 40% sequence identity) overlapped between the two sets. Subsequently, for binary classification, experimentally identified cleavage sites were labeled as positives, while all other non-cleavable residues in the same proteins were considered as negatives. The resulting dataset had 82 positive and 26,762 negatives, and we employed stratified sampling during model training to account for this class imbalanceness.

##### Conventional Sequence-based Features

We adopted four types of features proposed by [6]: 1) amino acid composition (AAC), 2) binary encoding (BE), 3) position-specific scoring matrix (PSSM), and 4) composition of K-spaced amino acid pairs (CKSAAP). See [6] for details of these features.

##### Experimental Setting

We evaluated three types of feature sets: **Baseline** (conventional sequence-based features only), **AF2** (the single representation (features) extracted from AF2) and **Extended** (**Baseline** + **AF2**).

To assess the discriminative power of these feature sets, we first employed representative machine learning (ML) classifiers and then complex deep learning (DL) models. See below for the detailed settings. Machine learning models were implemented using scikit-learn in Python, and deep learning models were built and trained using PyTorch.

Model performances were assessed using stratified 5-fold cross-validation with hyperparameter optimization (by grid search), followed by retraining on the full training set and evaluation on the independent test set. Performance was measured by the area under the ROC curve (AUC) and the area under the precision-recall curve (AUPR). The mean AUC (AUPR) over the five folds in 5-fold cross-validation was used as the final performance metric.

##### Machine Learning Model Setting

1. **Decision tree:** As an interpretable model, a decision tree (DT) classifier was used to capture direct hierarchical relationships between features and cleavage sites.
2. **Elastic net + kernel support vector machine:** We used a hybrid two-stage pipeline to achieve high performance in the high-dimensional space. First, elastic net (ElasticNet) was used as a feature selector to reduce the dimension and address multicollinearity by combining *L*_1_ and *L*_2_ regularization. Then, the selected features were input into a support vector machine (SVM) with a radial basis function (RBF) kernel, starting with default parameters (*C* = 1.0, *γ* = ‘scale’) for optimization.

Note that traditional classifiers require, as the input, a fixed-length vector for a given sequence. When we use **AF2** (and **Extended**), which is a matrix for a given sequence, we applied a flattening operation to transform the AF2 feature matrix into a vector. That is, the original AF2 matrix of 31 *×* 384 was transformed into a one dimensional vector with the size of 11,904 (=31*×*384). To facilitate the model convergence, we applied Z-score normalization (*StandardScaler*) to scale features based on the statistics of the training set.

##### Deep Learning Model Setting

We designed a hybrid neural network, which we call **MLP+CNN**, with two branches, i.e. a multi-layered perceptron (MLP) branch and a convolutional neural network (CNN) branch.

For **Baseline**, this network allows the effective processing of the heterogeneous properties of conventional feature types, as follows:

1. **MLP branch:** A MLP was employed to process global features (AAC) and flattened local features (BE and PSSM).
2. **CNN branch:** A 1D CNN (Conv1D) was used to extract local patterns from high-dimensional sequence motifs (CKSAAP). The two branches were trained independently by the respective feature types.

For **AF2** and **Extended**, the CNN branch can accommodate the single representation from AF2, where the input matrix of (31*×*384) is treated as 31 amino acids and 384 input channels. This means that the convolution filter was slid over the given sequence, to detect local structural motifs.

#### 2. Nucleic-acid-binding Site Prediction

##### Benchmark Datasets

We used proteins of the dataset in [7], under the condition that a sequence length is less than 1,00. Table 1 shows the statistics of the datasets we used.

**Table 1.** Statistics of two datasets in nucleic-acid-binding site prediction.

| Dataset | #proteins | #residues | #positives | #negatives | P/N ratio |
| --- | --- | --- | --- | --- | --- |
| *DNA-binding* |  |  |  |  |  |
| Train | 573 | 144,791 | 14,540 | 130,251 | 0.112 |
| Test | 129 | 22,403 | 1,352 | 21,051 | 0.064 |
| *RNA-binding* |  |  |  |  |  |
| Train | 494 | 136,345 | 14,584 | 121,761 | 0.120 |
| Test | 117 | 20,313 | 1,114 | 19,199 | 0.058 |

##### Conventional Sequence-based Features

We used four types of features: 1) PSSM, 2) hidden Makove model (HMM), 3) dictionary of secondary structure in proteins (DSSP), and 4) following [7], so-called atom features (AF), which comprise atom-level physicochemical properties, such as mass, charge, and Van der Waals radius.

##### Experimental Setting

We first evaluated our proposed feature set **AF2**, i.e. the single representation (features) from AF2, comparing with PSSM and HMM. In particular, we examined not only each of the three feature sets independently but also their combinations. Next, we generated a conventional sequence-based feature set, **Baseline**, which combines PSSM, HMM, DSSP and AF, where feature types for each residue were aggregated by global average pooling (GAP) and normalized to [0, 1] with MinMax scaling. We then compared three feature sets: **Baseline, AF2** (the single representation from AF2) and **Extended** (**Baseline**+ **AF2**).

To examine the discriminative power of these feature sets, we first used two representative ML models, i.e. SVM and random forest (RF) classifiers. They are selected to check the consistency of our results with those provided by a previous study [7]. The AF2 representation has high dimensionality and complex spatial dependencies. We then adopted a graph neural network (GNN) approach to naturally model non-Euclidean protein structures, decode latent spaces without manual dimensionality reduction, and restore spatial context through neighbor aggregation. Machine learning models were implemented using scikit-learn in Python, and deep learning models were built and trained using PyTorch.

We first used two types of validation: 1) internal validation: stratified 5-fold cross-validation. and 2) independent testing: models are tested on a held-out set. In both cases, performance was measured by AUC and AUPR.

##### Machine Learning Model Setting

1. **SVM:** we used a RBF kernel and *C* = 1.0. Probability estimates were enabled.
2. **RF:** 100 decision trees were used with the default hyperparameters.

##### Deep Learning Model Setting: Hierarchical Graph Neural Networks

Following [7], we utilized a hierarchical graph neural network (HGNN) to process the input, which can be defined as so-called attribute graph *G* = (*V, E*, **u**), where *V* is a node (amino acid residue) set, an edge in edge set *E* is generated by a spatial distance threshold (e.g., 14 Å) based on AF2-predicted coordinates and **u** is a set of feature vectors, each being assigned to a residue.

The initial input is a graph (with feature vectors) corresponding to a target protein, and after updating the feature vectors by information flow through edge connections, the probability of whether each residue serves as a nucleic acid-binding site or not is predicted through the architecture of HGNN (see [7] for the details of the architecture and learning algorithm of HGNN).

## Results

### 1. Calpain Cleavage Site Prediction

#### Performance Comparison

Table 2 shows the performance comparison among three feature sets, **Baseline, AF2**, and **Extended**, by using three different learning strategies. First, **AF2** outperformed **Baseline**, in five out of all six cases, implying that structural AF2 representation has richer discriminative power than simple sequence profiles. Second, **Extended** achieved the highest performances in five out of all six cases. More specifically, ElasticNet+kernel SVM with **Extended** achieved the best performance with AUC of 0.891 and AUPR of 0.075. In particular, **AF2** improved **Baseline**, and **Extended** further improved **AF2**, again implying the effectiveness of using AF2 single representation. Interestingly, this model clearly outperformed CNN+MLP both in AUC and AUPR, meaning that a concise, regularized model is more appropriate for this task than a complex, richer DL model. Additionally, we note that the performance level of **Baseline** was consistent with that of [6], showing the validity of the results in Table 2.

**Table 2.**
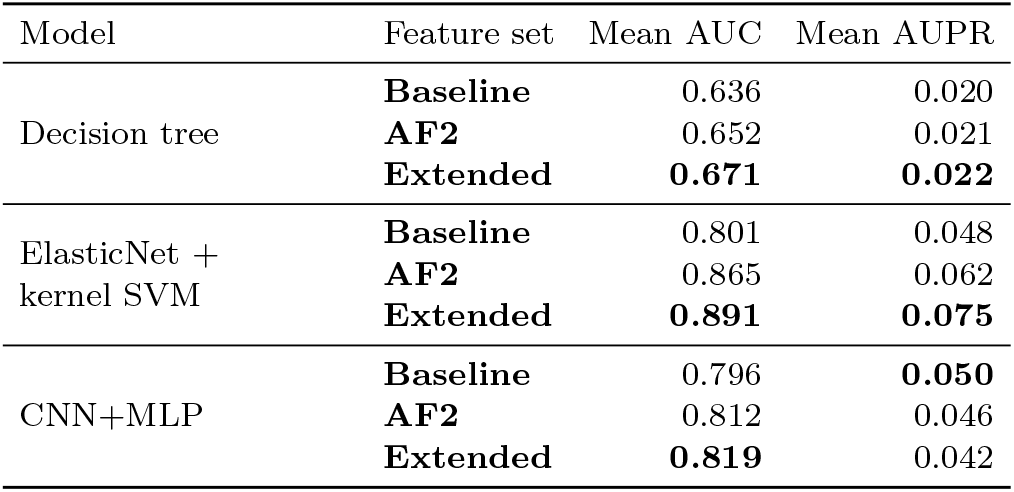
Performance comparison among three feature sets: Baseline, AF2 and Extended. Best results are bolded.

#### Feature Importance

For further feature analysis, we applied “SHapley Additive exPlanations” (SHAP) [9] to ElasticNet+kernel SVM, which achieved the best performance among the three learning strategies. Figure 2 shows the two beeswarm plots obtained when we used (A) **Baselilne** and (B) **Extended**. In (A), obviously, all top 20 crucial features are conventional sequence-based features, particularly 15 out of 20 coming from CKSAAP. On the other hand, in (B), all 20 top features are from single representation of AF2 and totally not from conventional sequence-based features, indicating that AF2 single representation was more important and useful in classification than conventional features.

**Figure 1:**
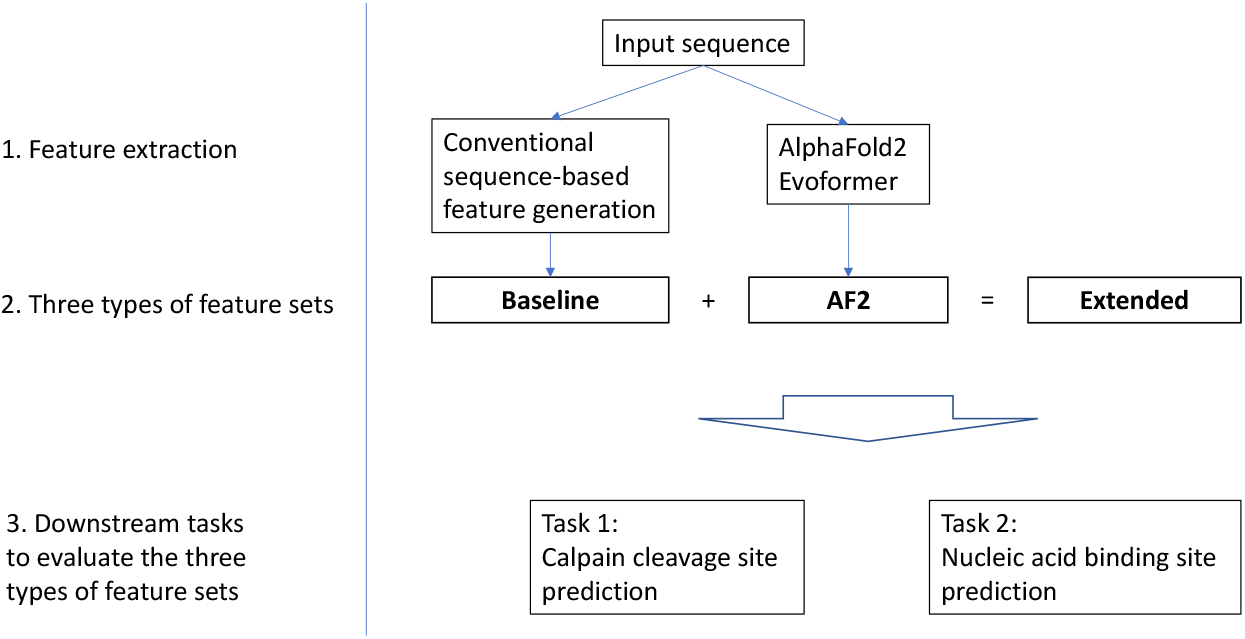
Overall framework of this work. Starting with the input sequence, we extracted features from the AlphaFold2 Evoformer, which were evaluated through two downstream tasks. comparing with conventional sequence-based features.

**Figure 2:**
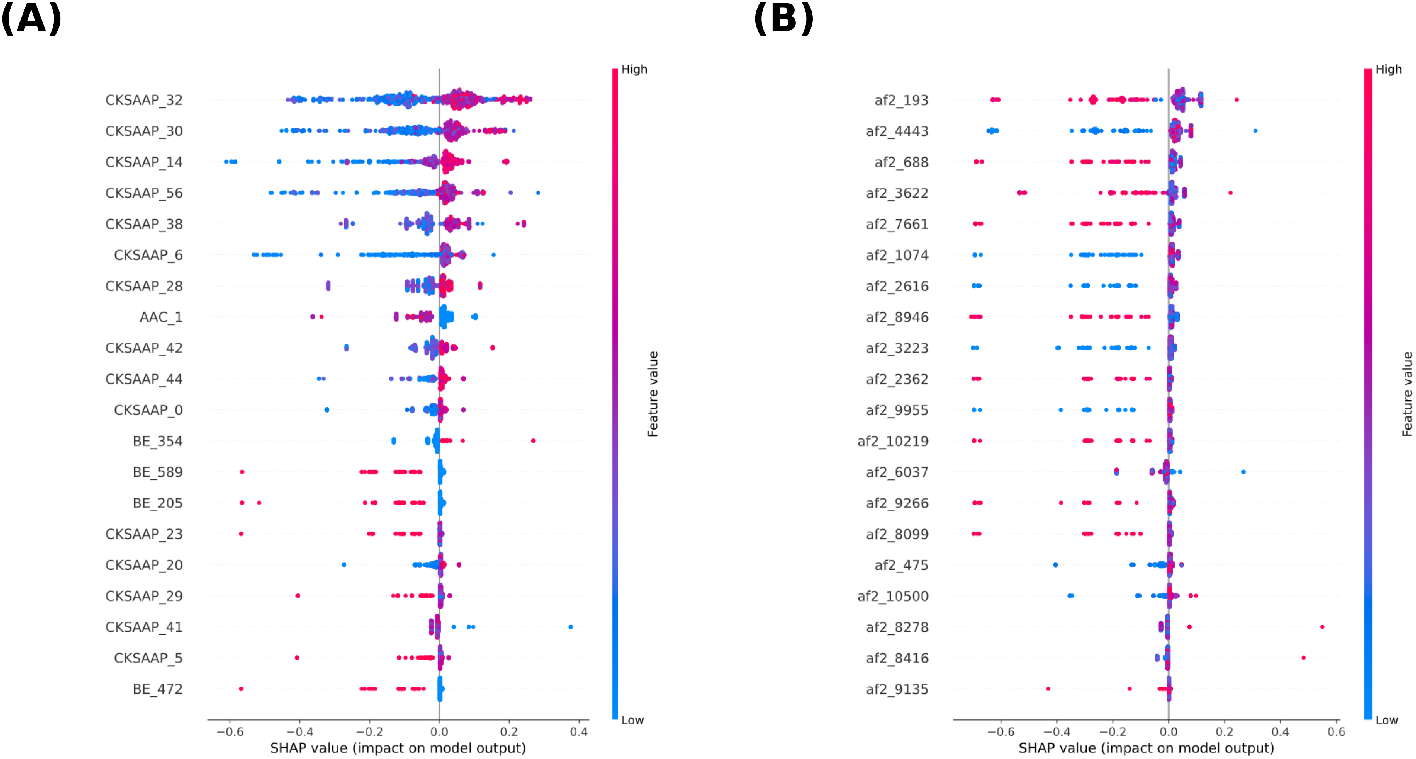
Results obtainecd by applying “SHapley Additive exPlanations” SHAP to ElasticNet+kernel SVM with: A) **Baseline** and B) **Extended**.

### 2. Nucleic-acid-binding Site Prediction

#### Machine Learning Model Performance Comparison

Table 3 shows the performance results of SVM and RF by using each of the three feature sets: PSSM, HMM and **AF2**, and their combinations. First, when we used SVM, **AF2** (AUC of 0.816) clearly outperformed PSSM (AUC of 0.693) and HMM (AUC of 0.629). This is also the case with AUPR. SVM with all features, i.e. PSSM + HMM + **AF2**, achieved the highest AUC, 0.817 (AUPR of 0.257). On the other hand, for RF, **AF2** was not necessarily the highest, however the highest performance by RF was AUC of only 0.729 (with PSSM + HMM) and AUPR of 0.173 (with HMM + **AF2**), still indicating that AF2 single representation can improve the performance due to the richness of structural information, under a proper learning model setting.

**Table 3.**
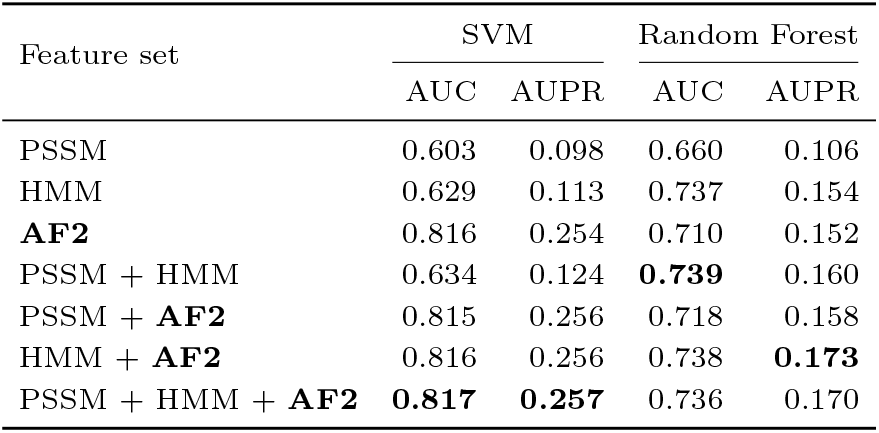
Performance comparison of different features, PSSM, HMM and AF2, and combinations when using SVM and RF.

#### Deep Learning Model Performance Comparison

Due to the high computational complexity of the DL model, we could not examine the features with the entire datasets. Instead, we focused on a specific part of the entire datasets, i.e. one test set, DNA-129 (129 DNA binding proteins) for DNA binding site and one test set, RNA-117 (117 RNA binding proteins) for RNA-binding site. They both have been often used as the most standard test sets for model evaluation in the literature, for example [7]. Table 4 shows the statistics of these two test sets.

**Table 4.** Statistics of DNA-129 and RNA-117, and that of “Easy Set” and “Hard Set”, which are both derived from RNA-117.

| Dataset | #proteins | #positives | #negatives |
| --- | --- | --- | --- |
| DNA-129 | 129 | 1,352 | 21,051 |
| RNA-117 | 117 | 1,114 | 19,199 |
| Easy Set | 11 | 105 | 1,805 |
| Hard Set | 106 | 1,009 | 17,394 |

Table 5 shows the AUC and AUPR for these two test sets, obtained by using three feature sets, **Baseline, AF2** and **Extended**. First, **AF2** outperformed **Baseline** in both cases. Second, **Extended** was better than **AF2** in one of the two cases. Overall, using **AF2** could enhance the classification performance of conventional sequence-based features.

**Table 5.** Performance comparison on the two independent test sets, where best results are in bold.

| Dataset | Feature Set | AUC | AUPR |
| --- | --- | --- | --- |
| DNA-129 | **Baseline** | 0.930 | 0.496 |
|  | **AF2** | 0.935 | 0.524 |
|  | **Extended** | **0.941** | **0.556** |
| RNA-117 | **Baseline** | 0.861 | 0.319 |
|  | **AF2** | **0.886** | **0.372** |
|  | **Extended** | 0.882 | 0.367 |

#### Performance Analysis with Changing Sequence Homology

For further analysis, we divided the test set into two subsets, according to the sequence homology: “Easy Set” (*>* 40% sequence identity to one or more sequences in the training set) and “Hard Set” (*<* 40% identity to any sequence of the training set). Table 4 shows the statistics of these two subsets. Figure 3 shows the AUC and AUPR obtained by applying SVM to the two subsets. First, from the results by Easy Set, **Baseline** outperformed **AF2** in both AUC and AUPR (for example, in AUC, 0.8620 by **Baseline** and 0.8542 by **AF2**). **Extended** further outperformed **Baseline** in both AUC and AUPR (for example, AUC of 0.8834 by **Extended**), meaning that **AF2**, which resulted in the worst performance among the three feature sets, could improve the performance of **Baseline** to reach that of **Extended**. This implies that **Baseline** and **AF2** have distinct, complementary information each other, for sequences homologous to the training data.

**Figure 3:**
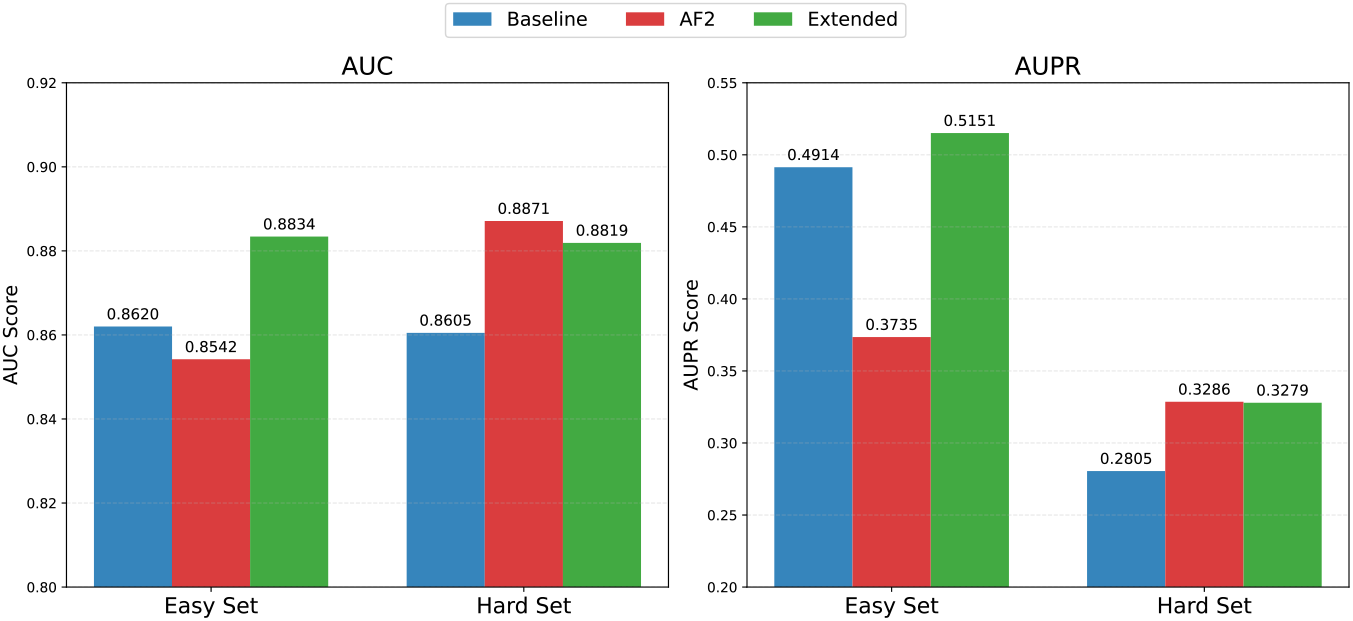
Performance comparison among three feature sets for two different homology levels between training and test sets, i.e. Easy Set (*>*40% of homology) and Hard Set (*<*40% of homology).

On the other hand, from the results of Hard Set, **AF2** outperformed **Baseline** in both AUC and AUPR (for example, in AUC, 0.8605 by **Baseline** and 0.8871 by **AF2**), indicating that AF2 single representation was more powerful than conventional sequence-based features. Interestingly, for Hard Set, the performance by **AF2** was unable to be beaten by **Extended** in both AUC and AUPR (for example, in AUC, 0.8819 by **Extended**). This means that for Hard Set, the information from **Baseline** was not much helpful and not necessarily complementary to that of **AF2. AF2**, i.e. AF2 single representation, could dominate the conventional sequence features, particularly for predicting novel, unknown proteins.

## Discussion

### Saliency Map to Examine the Importance of Surrounding Residues

Figure 4 shows so-called saliency visualization, generated by a Python library called Matplotlib, for the results of a specific part (Seq 5LJ3 G) of the RNA binding site data, obtained by HGNN, using (A) conventional sequence-based and (B) AF2 single representation. In each of the two figures, the graph in the 3D space shows that the connections from an edge-concentrated node, which corresponds to the focused residue, to the nodes corresponding to neighboring residues (in the 3D space). The color indicates the ‘gradient magnitude’ of each residue which means the importance of the corresponding residue for the focused residue (red to blue: more important to less important). From the result, neighboring residues are more blue colored in (B) than those in (A), indicating that neighboring residues are not so important in (B) than (A). That is, the information from neighboring residues is important when conventional sequence-based features are used, while this type of information is not so by AF2, implying that AF2 could capture more long-range interactions in the 3D space than neighboring information dependent sequence-based features.

**Figure 4:**
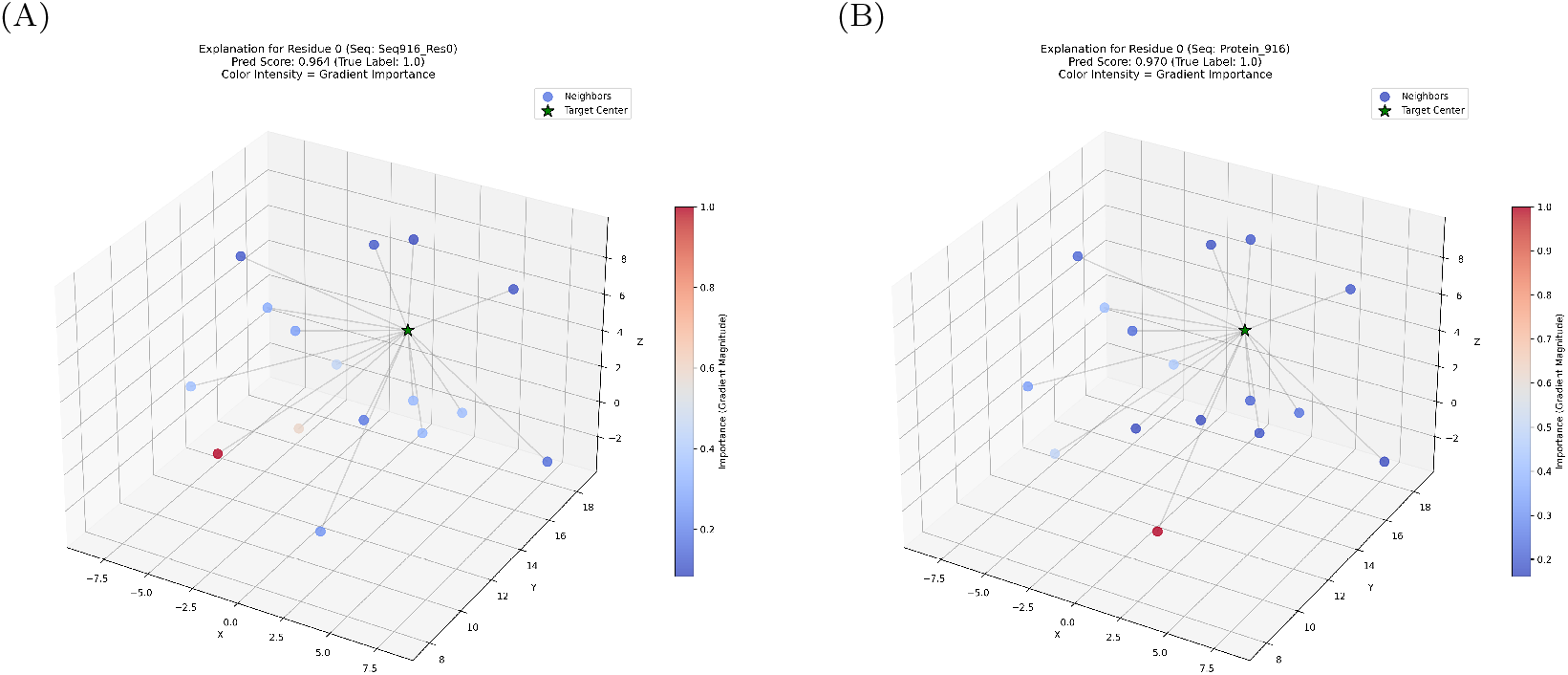
Saliency visualization comparison between two results obtained by HGNN with (A) conventional sequence-based feature sets and (B) **AF2** single representation.

This might be consistent with the result that SVM overwhelmed RF, where SVM could capture more long-range interactions than RF, due to the non-linear nature of SVM.

### Manifold Visualization by Dimension Reduction

For further analysis on the discriminative power of AF2 single representation, we projected the high-dimensional embeddings obtained by HGNN into a 2D space using Uniform Manifold Approximation and Projection (UMAP) [10]. Figure 5 shows the UMAP obtained by using (A) sequence-based features and (B) AF2, both extracted from RNA-117, where red and blue points are positives and negatives, respectively. To enhance the visual clarification, we randomly selected maximally 20,000 samples (points) per class. Red colored points might look more randomly distributed in (A) than in (B), implying that AF2 single representation might have generated more structured embeddings than conventional sequence-based features. This might have been a key to higher predictive performances by AF2 single representation.

**Figure 5:**
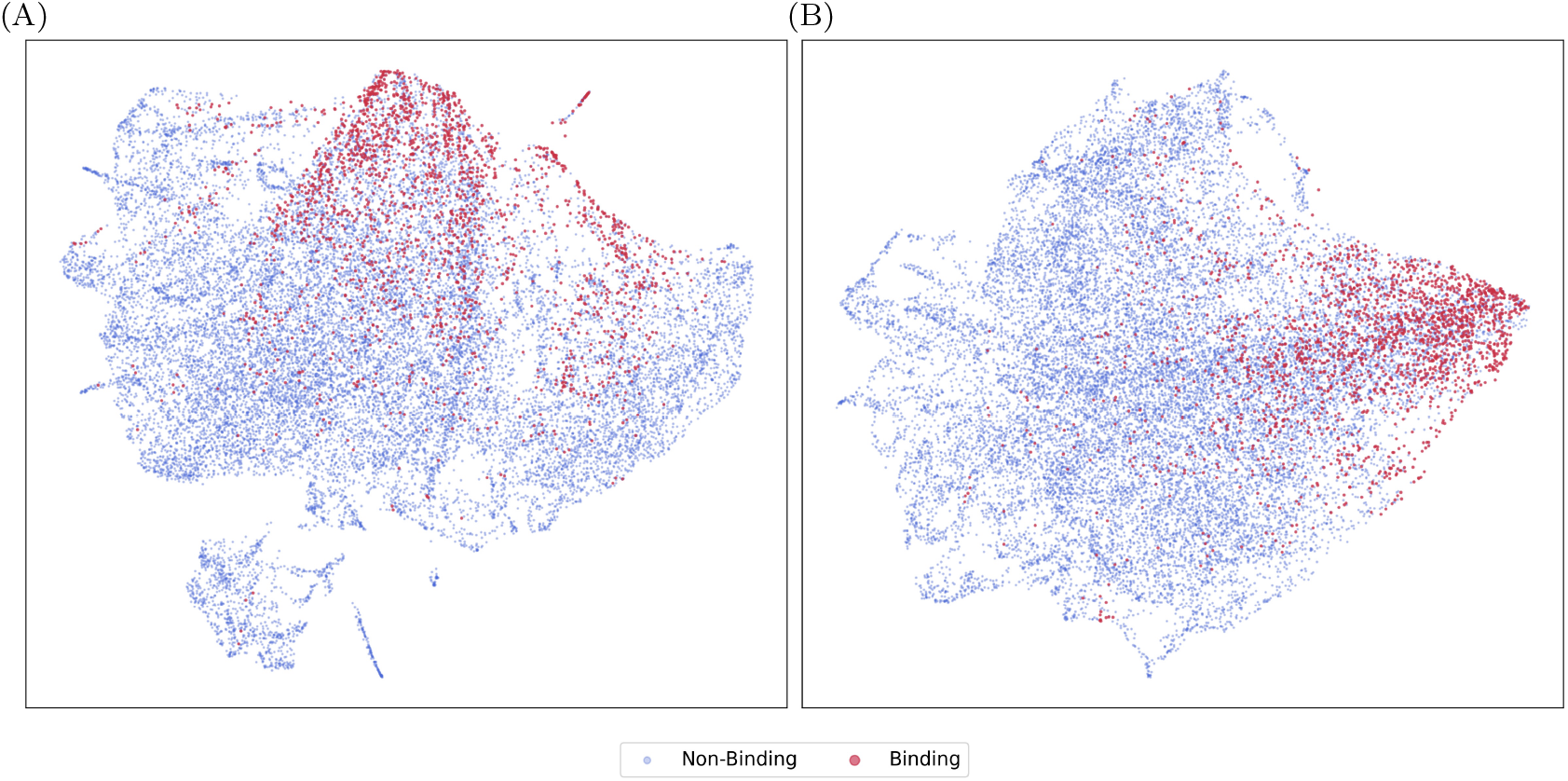
UMAP visualization comparison between two results for RNA-117, obtained by HGNN with (A) conventional sequence-based features and (B) **AF2**.

### Limitations

Despite the demonstrated advantages of AF2 single representation in sequence prediction tasks, we can raise at least three limitations of this representation: 1) computational loads: although bypassing the structure module by setting *num recycle=0* saves inference time notably, extracting single representation from the Evoformer needs higher computational load and more memory-intensive than generating standard sequence-based profiles. 2) static representation: many biological mechanisms, including calpain cleavage and nucleic acid site binding, have intrinsically disordered regions or require conformational shifts such as an induced fit. On the other hand, AF2 outputs a static, averaged conformational state. The current static features, AF2, inherently lack the capacity to capture such crucially temporal dynamics. 3) interpretability: while SHAP analysis can successfully identify important features, such as *af2 193*, mapping these specific latent features back to comprehensible physical or chemical properties, like charge, hydrophobicity, etc. remains an open challenge in representation learning.

## CONCLUSION

We have systematically assessed the effectiveness of applying extracted AlphaFold2 (AF2) single representation to biological sequence prediction tasks. In the two representative tasks, i.e. calpain cleavage and nucleic-acid binding site prediction, AF2 single representation consistently outperformed traditional sequence-based features. These tasks are highly imbalanced, and AF2 was proved to be highly effective for improving the performance of such an imbalanced data setting, e.g. kernel SVM for calpain cleavage site prediction (best performance: AUC of 0.891 and AUPR of 0.075). Importantly, the analysis using test sets divided by homology to training data showed two findings: AF2 single representaion 1) can be complementary information to the conventional sequence-based features for predicting high-homologous sequences, and 2) can dominate the sequence-based features for predicting low-homologous sequences. Particularly, the second point makes the AF2 single representation highly promising to be applied for any sequence input prediction tasks in bioinformatics.

As well as quantitative metrics, our interpretable analysis could visualize more detail of the improvement by AF2. SHAP analysis showed specific structural features by AF2 which occupied the list of the most important feature ranking. UMAP visualizations implied more structured or clustered embeddings of AF2 than sequence-based features in the reduced two-dimensional (2D) space.

Finally, we raise possible future work: 1) our results suggest that the AF2 single representation might be used for more challenging tasks, such as predicting ABC transporters or glycoenzymes, where, for precise prediction, more subtle, dynamic and region-specific embeddings (corresponding to 3D pockets) would be a key, which however simpler sequence-based features have struggled to identify. 2) we might be able to explore generating further computationally lighter feature extractors to address current computational issues for generating AF2 single representation. 3) recently, AlphaFold3 (AF3) [11] allows to predict multi-chain molecular interactions and complexes. Our idea in this work is, for an individual protein, to extract structure-aware feature representation, generally applicable to any downstream task, with which the target protein is involved. So our setting is totally different from that of AF3, meaning that the AF2 single representation would still remain useful, powerful and advantageous over traditional sequence-based features. Possible future work, however, would be to explore extracting similar intermediate representation from AF3, to capture latent embeddings, structurely representing, in allosteric contexts, multimolecular interactions, such as protein-protein, protein-ligand or protein-nucleic acid binding.

